# Effect of aging, endurance training, and denervation on innate immune signaling in skeletal muscle

**DOI:** 10.1101/2025.01.16.633423

**Authors:** Priyanka Khemraj, Anastasiya Kuznyetsova, David A. Hood

**Affiliations:** Muscle Health Research Centre, School of Kinesiology and Health Science, York University, Toronto, Ontario, M3J 1P3, Canada

**Keywords:** mitochondria, aging, denervation, endurance training, NLRP3 Inflammasome

## Abstract

Skeletal muscle function relies on mitochondria for energy and for mediating its unique adaptive plasticity. The NLRP3 inflammasome complex is an innate immune mechanism that responds to mitochondrial damage-associated molecular patterns (DAMPS), however its activity relative to mitochondrial dysfunction in muscle requires exploration. The purpose of this study was to characterize immune signaling and mitochondrial function in muscle during aging, endurance training, and disuse induced by denervation. Denervation led to decreases in muscle mass, mitochondrial content, and impaired respiration. Protein analyses revealed increases in NF-κB p65 and downstream inflammatory markers including NLRP3, caspase-1, GSDMD-N, STING and IL-1β, along with pro-apoptotic BAX and AIF. When assessing potential DAMPS, denervation led to increased ROS production but no changes in cytosolic mtDNA levels, relative to total mtDNA. Since we hypothesized that inflammasome activation would be increased with age, we studied young (6-8 months) and aged (21-22 months) mice that remained sedentary or underwent a 6-week voluntary running protocol. Aging resulted in marked increases in the expression of multiple pro-inflammatory and pro-apoptotic proteins. Remarkably, training uniformly attenuated age-related increases in BAX, NLRP3, caspase-1, STING, and GSDMD protein expression, and reduced the elevated level of cytosolic mtDNA evident in aged muscle. Training adaptations were evident also in the aged animals by the preservation of muscle mass and improvements in oxygen consumption and endurance performance and were achieved despite a lower training distance than in young animals. Our results strongly implicate endurance training as a promising therapeutic for combatting disuse and age-related inflammation in skeletal muscle.

## Introduction

The innate immune system is a highly reactive network that aids in the recognition of, and response to harmful stimuli, ultimately preserving cellular homeostasis (1). Additionally, the immune system promotes skeletal muscle maintenance by mediating muscle regeneration, pathogenic invasion, and short-term muscle damage to retain optimal tissue function (1, 2). However, while this response is beneficial acutely, chronic inflammation can augment deteriorations in skeletal muscle health and induce atrophy (3, 4). As a component of the innate immune system, the NLRP3 inflammasome complex can respond to a variety of pathogen-associated molecular patterns (PAMPs) and endogenous damage-associated molecular patterns (DAMPs) including viral RNA and ion fluctuations (5). To regulate its activation, the NLRP3 inflammasome complex uses a two-step activation process. The priming stage is initiated by the recognition of DAMPs and PAMPs by pattern recognition receptors such as Toll-like receptors and other sensors such as the cGAS-STING pathway (6, 7). This promotes the nuclear translocation of NF-κB, which aids in the transcriptional upregulation of NLRP3, pro-IL-1β and pro-IL-1. Following this step, a second signal initiates the assembly of the multi-protein complex comprised of NLRP3, apoptosis speck-like protein (ASC), and the cysteine protease procaspase-1. The subsequent formation of the complete NLRP3 inflammasome complex leads to the maturation of IL-1β, IL-18, and pyroptosis initiator gasdermin-D to aid in the destruction of dysfunctional cells and mount a localized immune response (5, 8). Recently, there has been an increased focus on investigating how mitochondrial adaptations can influence NLRP3 inflammasome activation and skeletal muscle health. Due to their roles as signaling hubs, skeletal muscle mitochondria have been implicated in the release of several DAMPS including ROS, mitochondrial DNA (mtDNA), and cardiolipin that can trigger an innate immune response (9). Chronic skeletal muscle disuse brought about by nerve transection is associated with increased mitochondrial dysfunction, muscle atrophy and reduced oxidative capacity (10). As a result of a fragmented mitochondrial reticulum and the accumulation of mtDAMPS such as mtROS and mtDNA, prior literature has observed a pro-inflammatory profile within skeletal muscle following periods of chronic disuse that is augmented by increased NLRP3 inflammasome activity and has been shown to contribute to skeletal muscle wasting (11, 12). In addition, the process of aging is also associated with similar adaptations including impairments in several mitochondrial quality control processes, leading to a less functional mitochondrial pool and a chronic state of low-grade inflammation which has been coined “inflammaging” (13). The NLRP3 inflammasome complex has been implicated in contributing to this pro-inflammatory environment within skeletal muscle, as overactivation of caspase-1 facilitated increased muscle atrophy alongside reductions in force and endurance capacity, which was rescued upon knockout of NLRP3 (14). In contrast, chronic exercise has been widely considered one of the best interventions for skeletal muscle by promoting an interconnected mitochondrial network that is characterized by increased mitochondrial content and respiration and the reduced production of damage molecules such as ROS (15). In other tissue types, long-term aerobic training has been associated with a reduced inflammatory environment and a decreased drive toward NLRP3 inflammasome activation (16, 17). However, the relationship between chronic exercise, mitochondrial health and NLRP3 inflammasome activation in muscle has yet to be explored.

These findings have prompted us to further investigate the relationship between mitochondrial adaptations and NLRP3 inflammasome activation *in vivo* using mouse models of chronic muscle disuse and training to elucidate the impact of these opposing stimuli. In this study, we induced muscle disuse via unilateral hindlimb denervation in young mice and measured mitochondrial function and NLRP3 inflammasome activation. In addition, we studied the impact of endurance training on inflammasome activation using both young and aged mice. Our findings indicate a positive relationship between skeletal muscle disuse, maladaptive mitochondrial adaptations, and NLRP3 inflammasome activation leading to a pro-inflammatory environment. In contrast, chronic exercise led to improvements in skeletal muscle mitochondria and the attenuation of innate immune signaling in aged mice. Thus, these findings highlight a relationship between mitochondrial function and inflammasome activation that is therapeutically modifiable by exercise.

## Methods

### Animals

All mice were obtained from Jackson Laboratories and housed in the York University vivarium with a 12:12h light-dark cycle and open access to food and water. Approval for all experiments was obtained from the York University Animal Care Committee. For the denervation study, male FVB mice were 3-6 months at the time of tissue collection. For the voluntary wheel running study, young mice were 6-8 months old and aged mice were 21-22 months old at the time of tissue collection.

### Sciatic Transection Surgery

At 3-6 months of age, male wildtype FVB mice were subjected to a sciatic nerve transection to induce hindlimb muscle denervation. Briefly, mice were anesthetized using isoflurane, and a small incision was made to reveal the sciatic nerve. In the denervated limb, a 2-3mm section of the nerve was transected and the incision was subsequently close using sutures and staples. The contralateral limb was sham-operated and used as an internal control. For post-surgery pain management, mice were given subcutaneous injections of Metacam for two days post-surgery (Day 1: 2µg/g body weight; Day 2: 1µg/g body weight). Additionally, all mice were also provided a 25mg/100mL dose of Baytril within their water to reduce the risk of post-surgery infection for the duration of the week. Seven days following denervation, hindlimb muscles were collected and flash-frozen in liquid nitrogen for subsequent analysis.

### Voluntary Wheel Running

Male young (4-5 months) and old (19-20 months) C57BL6/J mice were randomly assigned to either a sedentary control group or a training group, so that they were approximately 6-8 months and 21-22 months at the time of tissue collection. Mice in the training group had their cages fixed with a freely rotating running wheel for the remainder of the training intervention. Animals were given a one-week acclimatization period followed by a 6-week training period where revolutions were recorded by a magnetic counter. Distance in kilometres was calculated by multiplying the circumference of the wheel by the number of revolutions run.

### Acute Endurance Testing Protocol

An acute endurance test was performed for all animals before and after the sedentary or training period using a mouse treadmill. Mice were acclimatized to the treadmills for two consecutive days before testing (Day 1: 0 m/min for 5 minutes, 5 m/min for 5 mins, 10 m/min for 10 minutes; Day 2: 0 m/min for 5 minutes, 5 m/min for 5 minutes, and 10 m/min for 10 minutes). To avoid injury to the aged animals, the young and aged animals were tested with a less vigorous treadmill protocol. Exhaustion was defined as the inability to remain on the treadmill despite prodding. Total distance and time ran were used to calculate percent changes as a measure of improvement.

### Muscle protein extraction

Protein extracts from the tibialis anterior muscle were prepared from 25-30g of flash-frozen mouse muscle. The tissue was immersed in a Sakamoto Extraction Buffer (20mM HEPES, 2mM EGTA, 1% Triton X-100, 50% Glycerol, 50mM B-Glycerophosphate) containing protease and phosphatase inhibitors (Sigma) to prevent protein degradation. The tissue was then mechanically homogenized using a Tissue Lyser (Qiagen) at the frequency of 30Hz in one-minute intervals until a uniform homogenate was obtained. To isolate the protein extract, the homogenized tissue was centrifuged at 14000g for 10 minutes at 4°C, and the resulting supernate was stored at -80°C until further analysis.

### Mitochondrial and Cytosolic Fractionations

Fractionation was performed on fresh gastrocnemius tissue. Immediately following collection, muscle was immersed a Mitochondrial Isolation Buffer. Muscle was minced on a glass plate over ice and subsequently homogenized using a dounce homogenizer (Fisher Scientifics). The resulting homogenate was centrifuged at 1200g for 15 minutes, and the resulting supernate was collected and centrifuged at 21000g for 20 minutes to obtain a mitochondrial pellet and cytosolic fraction. The mitochondrial pellet was resuspended in 100µL of the mitochondrial isolation buffer and centrifuged at 21000g for 20 minutes. The supernate was discarded and the final pellet was resuspended in 20-40µL of buffer to obtain the final mitochondrial fraction. The cytosolic fraction was centrifuged twice at 21000g, where the supernatant was transferred to a new Eppendorf tube each time to obtain the final cytosolic fraction. All centrifugation steps were performed at 4°C. Both the mitochondrial and cytosolic fractions were stored at -80°C until further analysis. Protein concentrations were assessed using a Bradford protein assay and purity was confirmed by western blotting using mitochondrial content markers, COX-IV and VDAC to ensure cytosolic purity.

### Western Blotting

Protein concentrations were determined using a Bradford Protein Concentration Assay and a plate reader. For western blotting, 15-90µg of protein was loaded onto a 10-15% SDS-page gel and separated by molecular weight using electrophoresis at 120 volts for approximately 90 minutes. The protein was then transferred to a nitrocellulose membrane and stained using Ponceau red to ensure proper transfer and for subsequent normalization. The membranes were then blocked with 5% skim milk in 1xTBS-T for 1 hour and incubated with the specific primary antibody overnight at 4°C. The following day, the blots were washed with TBS-T (5 minutes x 3 washes) and incubated with the corresponding secondary antibody for 1 hour at room temperature. All antibodies used in this study are listed in Table 1. The blots were washed again (5x3) and imaged using enhanced chemiluminescence via the iBright CL1500 Imaging System. All samples were normalized to the corresponding ponceau to correct for protein loading.

**Table 1:**
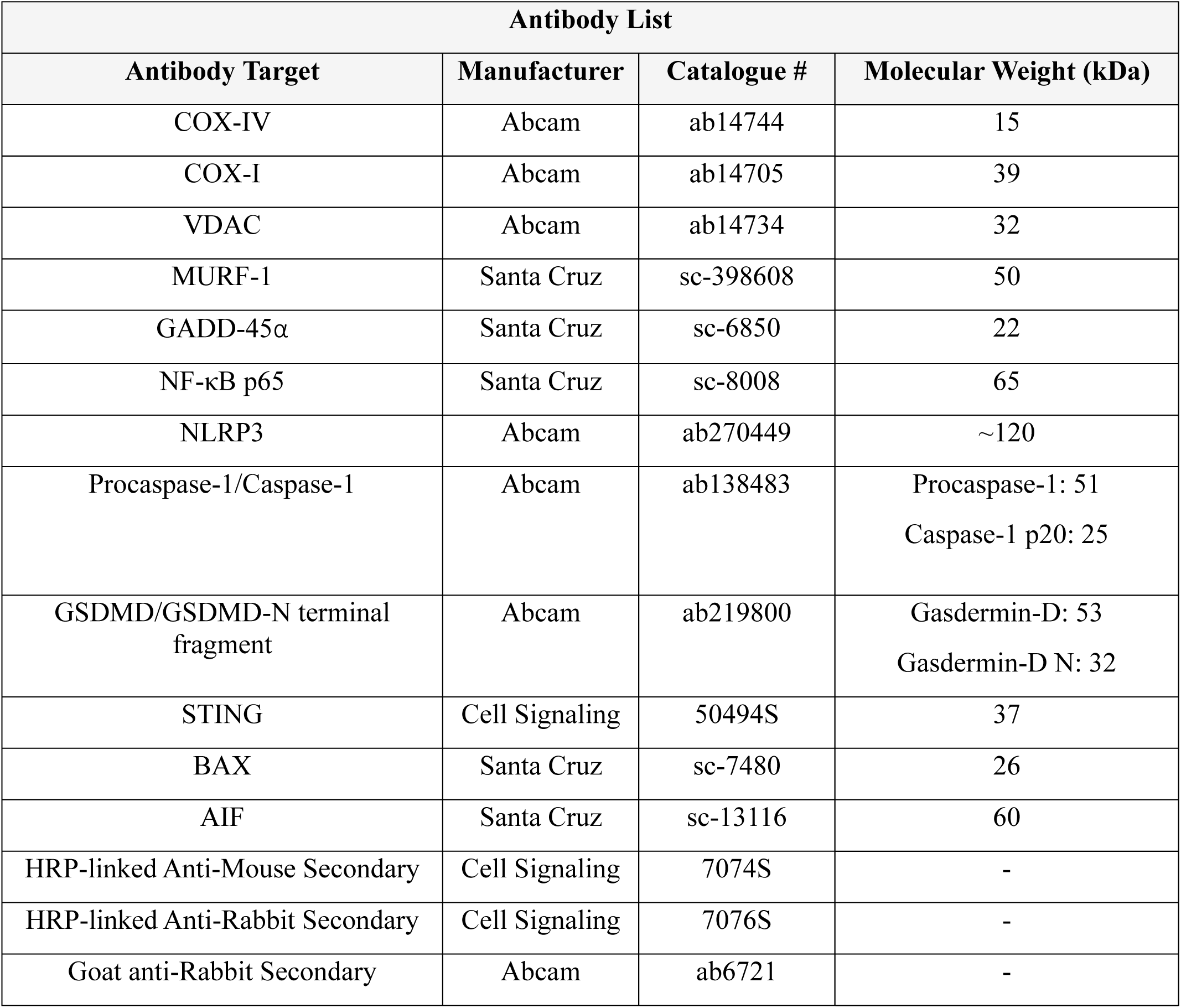
List of antibodies used for western blotting.

### ELISA

The Mouse IL-1β/IL-F2 Duoset ELISA kit (R&D Systems) was used to assess the protein levels of IL-1β. Within a 96-well microplate, the IL-1β capture antibody was added to each well and incubated overnight at room temperature. The following day, each well was washed and blocked with 1% BSA for one hour at room temperature. Following this incubation, the plate was washed, and the corresponding skeletal muscle extracts and protein standard (15-1000 pg/mL) was added and incubated for two hours at room temperature. Afterward, the IL-1β detection antibody was added and incubated for two hours at room temperature. The Streptavidin-HRP conjugate was then added and incubated, followed by the addition of a 1:1 hydrogen peroxide and tetramethyl benzadine substrate mixture for 20 minutes at room temperature. Finally, the stop solution was added to each well and the resulting absorbance was measured using a plate reader at 450nm and 570nm. To correct for optical inconsistencies and baseline absorbance, the 570nm values were subtracted from the 450nm measurements, and the blank solution absorbance was subtracted from the resulting values. To assess the concentration of IL-1β within the samples, the measured concentrations were compared to the standard curve.

### gDNA extraction

Genomic DNA was isolated from the gastrocnemius muscle and the cytosolic fraction for subsequent qPCR analysis using the DNeasy Blood and Tissue Kit (Qiagen) as per the manufacturer’s instructions. For the whole muscle extraction, ∼18-25 mg of frozen muscle was immersed in 180µL of Buffer ATL and 20µL of proteinase K and incubated at 56°C for 1 hour and 20 minutes. Afterward, 200µL of Buffer AL and 200µL of absolute ethanol were added, and the resulting mixture was added to the DNeasy spin column. The mixture was then centrifuged at 6000g for 1 minute, washed with 500µL of Buffer AW1, and spun again at 6000g for 1 minute. Following this spin, 500µL of Buffer AW2 was added and centrifuged at 20000g for 3 minutes. To elute the DNA, 120µL of Buffer AE was added for 1 minute, and centrifuged for 6000g for 1 minute. The resulting gDNA was collected and stored at -20°C. All centrifugation steps were performed at room temperature. For the cytosolic gDNA extraction, 200µL of the cytosolic fraction was combined with 20µL of proteinase K and 200µL of buffer AL, before being incubated at 56°C for 10 minutes. For the remaining steps, the same protocol was followed as for the gDNA extraction from whole tissue. DNA concentrations were measured using the NanoDrop 2000 and the cytosolic and gastrocnemius samples were diluted to equal concentrations, respectively. These concentrations underwent primer optimization before experimental qPCR plates were performed.

### Qualitative Polymerase Chain Reaction

To analyze genes of interest, forward and reverse primers were created and optimized to the corresponding set of samples (Table 2). mRNA expression is measured using the StepOnePlus Real-time PCR system (Applied Biosystems) along with the SYBR Green qPCR mix. Each well plate contains 25µL of reaction volume consisting of sterile water, the corresponding forward and reverse primers (20µM) and 2µL of gDNA. All samples are run in duplicates to increase reliability. The resulting gene expression is then normalized to the housekeeping gene, GAPDH, and quantified. Quantification is done by 1) Calculating the ΔCycle Threshold (CT) = mean CT of gene of interest – mean CT of housekeeping genes, 2) The resulting fold change is calculated by 2^(-ΔCT)^. The obtained whole muscle fold changes were corrected to account for the increased dilution factor in the whole muscle samples compared to the cytosolic fractions.

**Table 2:**
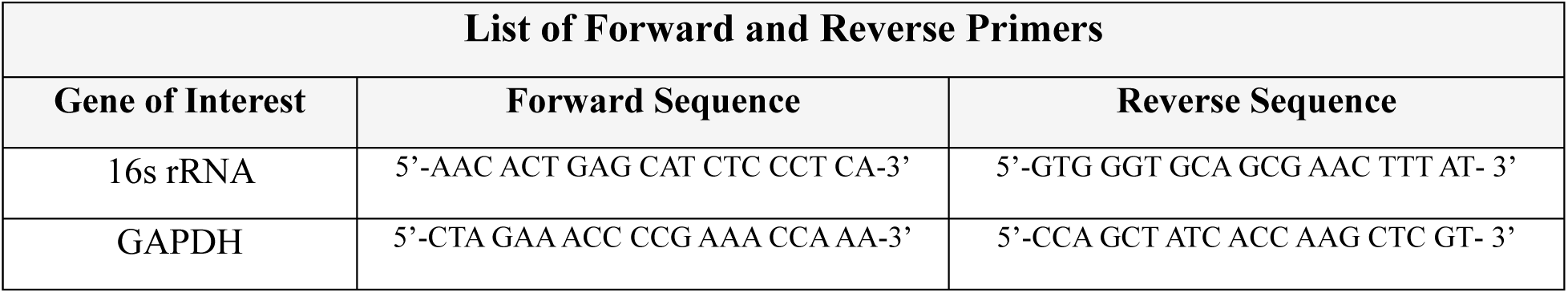
qPCR primers used in the analysis of mitochondrial DNA.

### High-resolution Respiratory and H_2_O_2_ Production

A small portion of the tibialis anterior muscle was collected upon tissue removal and stored a BIOPS buffer (2.77mM CaK_2_EGTA, 7.23mM K_2_EGTA, 5.77mM Na_2_ATP, 6.56mM MgCl_2_ hexahydrate, 20mM Taurine, 15mM Na_2_Phosphocreatine, 20mM Imidazole, 0.5mM Dithiothreitol, 50mM MES Hydrate). Immediately following collection, the muscle fibers were teased apart and permeabilized in a BIOPS buffer with 40µg/µL Saponin for 30 minutes at 4°C on an orbital shaker with constant motion. The fiber bundles were then washed twice in Buffer Z (105mM K-MES, 30mM KCl, 10mM KH_2_PO4, 5mM MgCl_2_ hexahydrate, 1mM EGTA, 5mg/mL BSA). Following this preparation, the fibers were incubated in Buffer Z with 1µM Blebbistatin to avoid tetanic contractions (B592500, Toronto Research Chemicals), 10µM Amplex-Red (A36006, Thermofisher), 25 U/mL Cu/Zn SOD1 to convert any superoxide molecules to H_2_O_2_, and 2mM EGTA. For ROS measurements, 0.5U horseradish peroxidase and 0.1µM H_2_O_2_ were added to the chamber for calibration. To obtain measurements for various states of respiration, high-resolution respirometry was used (Power Oxygraph 2K-Fluorospirometer, Oroboros Instruments) and substrates were added as designated: 1) Complex I inactive: pyruvate and malate, 2) Complex I active: ADP, and 3) Complex I+II active: succinate.

### Statistical Analysis

All data was analyzed using GraphPad Prism 9.0, and values were expressed as mean ± standard deviation. Differences between the denervated and contralateral limb was calculated using a student’s paired t-test. For the VWR study, all data was analyzed using a two-way ANOVA with a Tukey’s post-hoc test to assess differences between groups. Statistical significance was considered as p < 0.05.

## Results

### 7 days of muscle denervation alters skeletal muscle and mitochondrial functioning

To induce muscle disuse, a sciatic nerve transection was performed (Figure 1A). Following 7 days of muscle denervation, there was a drastic reduction in total hindlimb muscle mass (Figure 1C, p<0.0001), but no reduction in total body weight (Figure 1B). This effect can be seen consistently across skeletal muscles of various fiber types including mixed muscles such as the tibialis anterior (Figure 1D, p<0.0001), the gastrocnemius (Figure 1F, p<0.001), and the soleus (Figure 1E, p<0.0001). To further confirm the presence of muscle atrophy, the protein expression of several muscle atrophy markers was assessed. Muscle denervation induced a drastic increase of the E3 ubiquitin ligase, MURF1 (p<0.001), and muscle atrophy inducer, GADD45α (Figure 1G, p<0.05), demonstrating increased signaling towards muscle degradation.

**Figure 1:**
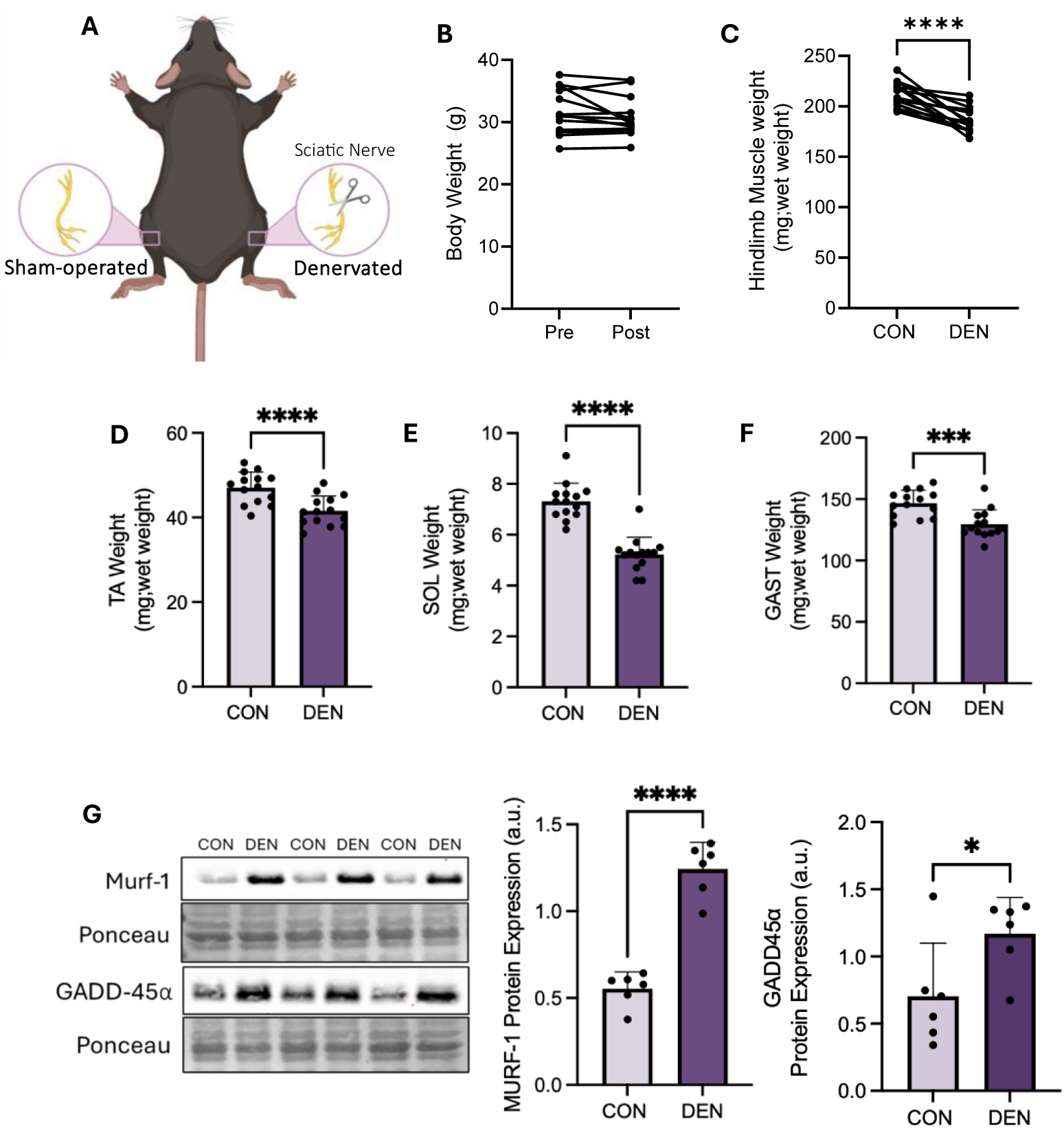
Skeletal muscle characteristics following 7 days of muscle denervation. Study design for the induction of skeletal muscle disuse using a sciatic nerve transection (A). Differences in body weight before and after denervation surgery (B) (n=14). Total hindlimb muscle mass (C) and individual muscle weights of TA (D), SOL (E), and (F) GAST compared to sham-operated control (n=14). Representative western blot of MURF-1 and GADD-45α protein expression alongside the corresponding quantifications (G) (n=6). All measurements underwent a paired student’s t-test. *, p<0.05; ***, p<0.001; ****, p<0.0001). TA, tibialis anterior; SOL, soleus; GAST, gastrocnemius; CON, control; DEN, denervated. Values are expressed as mean ± SD. A portion of Figure 1A was created using BioRender.com.

### Muscle denervation induces maladaptive changes within skeletal muscle mitochondria

Next, to assess the impact of denervation on the mitochondrial reticulum, we evaluated several mitochondrial markers including the outer membrane protein VDAC, and a subunit of complex IV within the electron transport chain, COX-IV. Both content markers demonstrated reduced protein expression following the sciatic nerve transection indicating reduced mitochondrial content compared to the sham-operated control (Figure 2A, B, p<0.05; Figure 2C, p<0.01). Furthermore, to examine the health of the mitochondrial pool, respiration analysis was performed using O_2_k high-resolution respirometry. Muscle disuse resulted in impaired complex I inactive (Figure 2D, p<0.05) and active respiration (Figure 2E, p=0.05) when supplemented with pyruvate/malate and ADP respectively, indicating maladaptive mitochondrial functioning.

**Figure 2:**
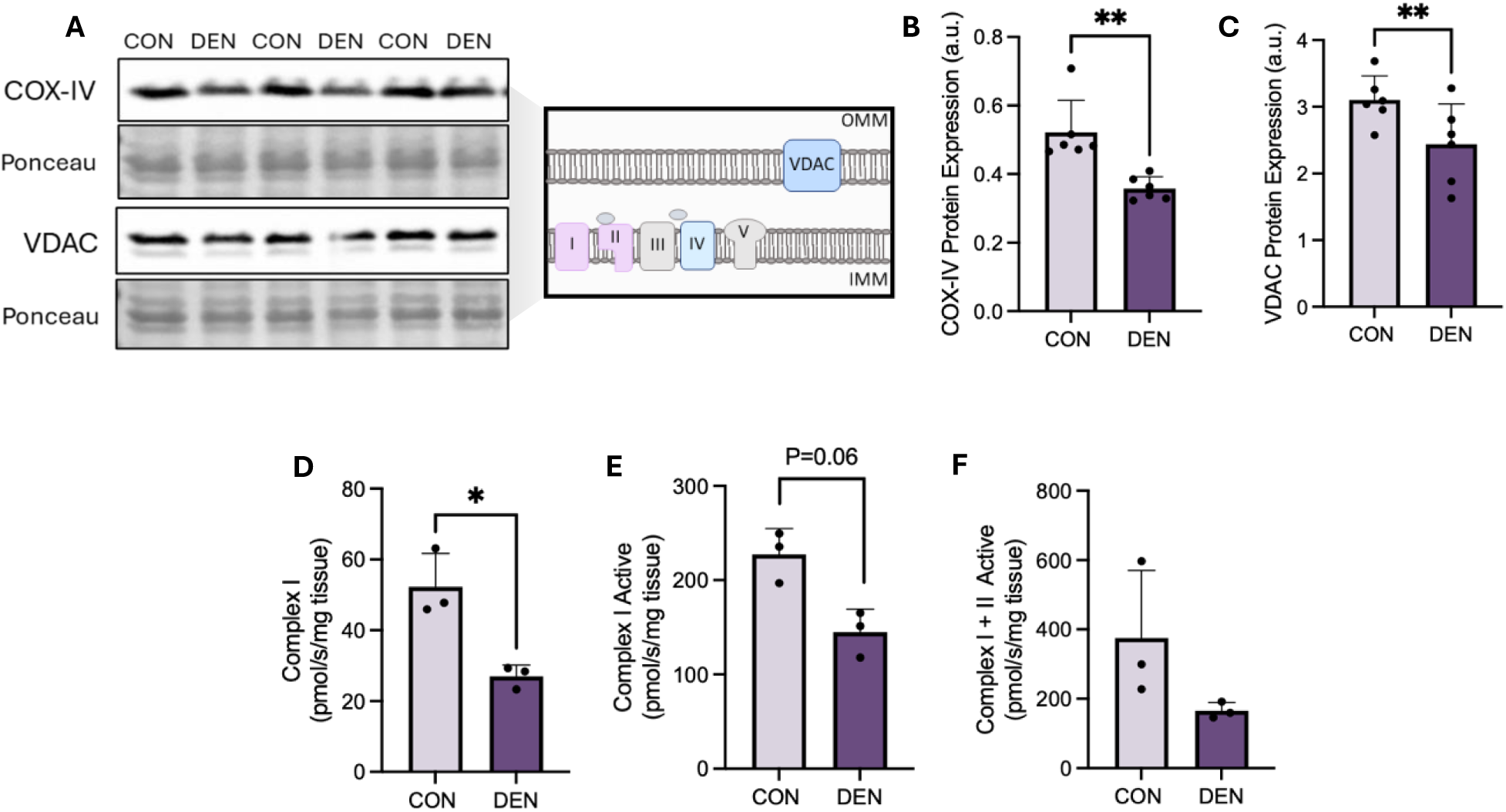
Skeletal muscle mitochondrial adaptations following 7 days of muscle denervation. Representative western blots of mitochondrial content markers, COX-IV and VDAC (A), alongside their corresponding quantifications (B, C) (n=6). Oxygen consumption rates during complex I (D), complex I active (E), and complex I+II active (F) states of respiration (n=3). Data was analyzed using a paired student’s t-test. *, p<0.05; **, p<0.01. CON, control; DEN, denervated; OMM, outer mitochondrial membrane; IMM, inner mitochondrial membrane. Values are expressed as mean ± SD.

### NLRP3 inflammasome activation is upregulated following 7 days of denervation

To elucidate the effect of denervation on NLRP3 inflammasome activation, the protein expression of various components of the signaling pathway was measured. Following denervation, there was a drastic increase in NF-κB subunit p65 expression, which partly drives the upstream transcriptional activation of the NLRP3 inflammasome complex (Figure 3A, 3B, p<0.00001). Next, to understand how the structural components of the NLRP3 inflammasome complex were affected, we measured the protein levels of NLRP3 and procaspase-1 protein. Notable increases in NLRP3 (Figure 3C, p<0.01) and procaspase-1 protein expression (Figure 3E, p<0.01) were observed. To characterize NLRP3 inflammasome activation within denervated skeletal muscle, downstream protein targets within the signaling pathway were measured. Muscle disuse led to an increased expression of caspase-1 p20 (Figure 3D, p<0.05), and its downstream targets GSDMD (Figure 3G, p<0.001) and its active N-terminal fragment, GSDMD-N (Figure 3F, p<0.05). Furthermore, disuse was associated with increases in the cytokine IL-1β, suggesting an increased pro-inflammatory environment within denervated hindlimb muscle (Figure 3H, p=0.05).

**Figure 3:**
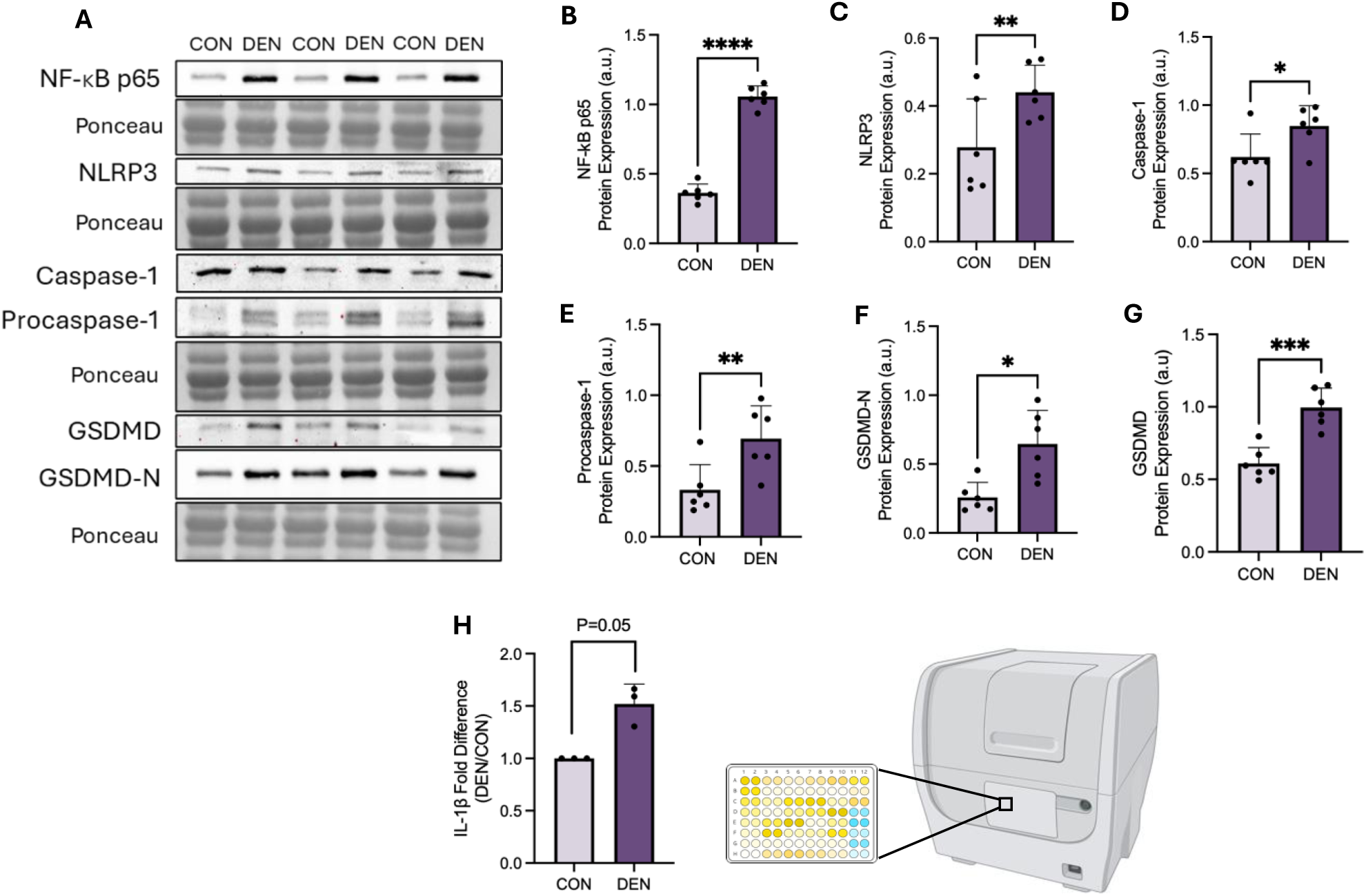
NLRP3 Inflammasome components following muscle denervation. Representative western blot of proteins involved in the NLRP3 Inflammasome signaling pathway (A), including NF-κB p65 (B), NLRP3 (C), Caspase-1 p20 (D), Procaspase-1 (E), GSDMD-N (F) and GSDMD (G) (n=6). Fold difference of IL-1β protein levels (DEN/CON) assessed by ELISA (H) (n=3). Data was analyzed using a paired student’s t-test. *, p<0.05; **, p<0.01; ***, p<0.001; ****, p<0.0001. CON, control; DEN, denervated; GSDMD, Gasdermin-D; GSDMD-N, Gasdermin-D N-terminal fragment. All values are expressed as mean ± SD. A portion of Figure 3H was created using BioRender.com.

### Muscle denervation is associated with alterations in DAMP release and increased mPTP pore protein expression

Since we established increased NLRP3 inflammasome activation in denervated muscle, we investigated the presence of DAMPS that could mediate the pro-inflammatory environment. Denervation induced remarkable increases in ROS compared to the basal levels present in the sham-operated control tissue (Figure 4A, 4B, 4C, p<0.05). Another potent DAMP is mtDNA which can be released into the cytosol from abnormally functioning mitochondria. As expected, the quantity of mtDNA within the whole gastrocnemius muscle was significantly decreased following denervation (Figure 4D, p<0.05), in line with the reduction in mitochondrial content observed (Fig. 2). As a result, mtDNA levels within the cytosolic fraction were also reduced (Figure 4E, p<0.05) and the proportion of mtDNA within the cytosol corrected to whole muscle mtDNA was not altered significantly following the 7-day denervation period (Figure 4F). We then investigated upstream and downstream pathways related to mtDNA release that are involved in NLRP3 inflammasome activation. The cGAS-STING DNA-sensing pathway is directly responsive to mtDNA, and approximate 3-fold increases were observed in both cGAS (Figure 4H, p=0.05), and STING (Figure 4I, p<0.0001) protein expression following denervation. This clearly suggests an increased drive towards the activation of cytosolic DNA sensing pathways within denervated muscle. In addition, since the opening of the mPTP has been previously implicated in augmenting the release of DAMPS to promote inflammation, we measured several proteins that are involved in this process. As seen in Figure 4J and 4K, denervation was associated with increases in the pro-apoptotic BAX and AIF protein when corrected to mitochondrial content (p<0.01, p<0.05, respectively), which illustrates the altered mitochondrial composition that accompanies denervation, and that promotes inflammasome activation.

**Figure 4:**
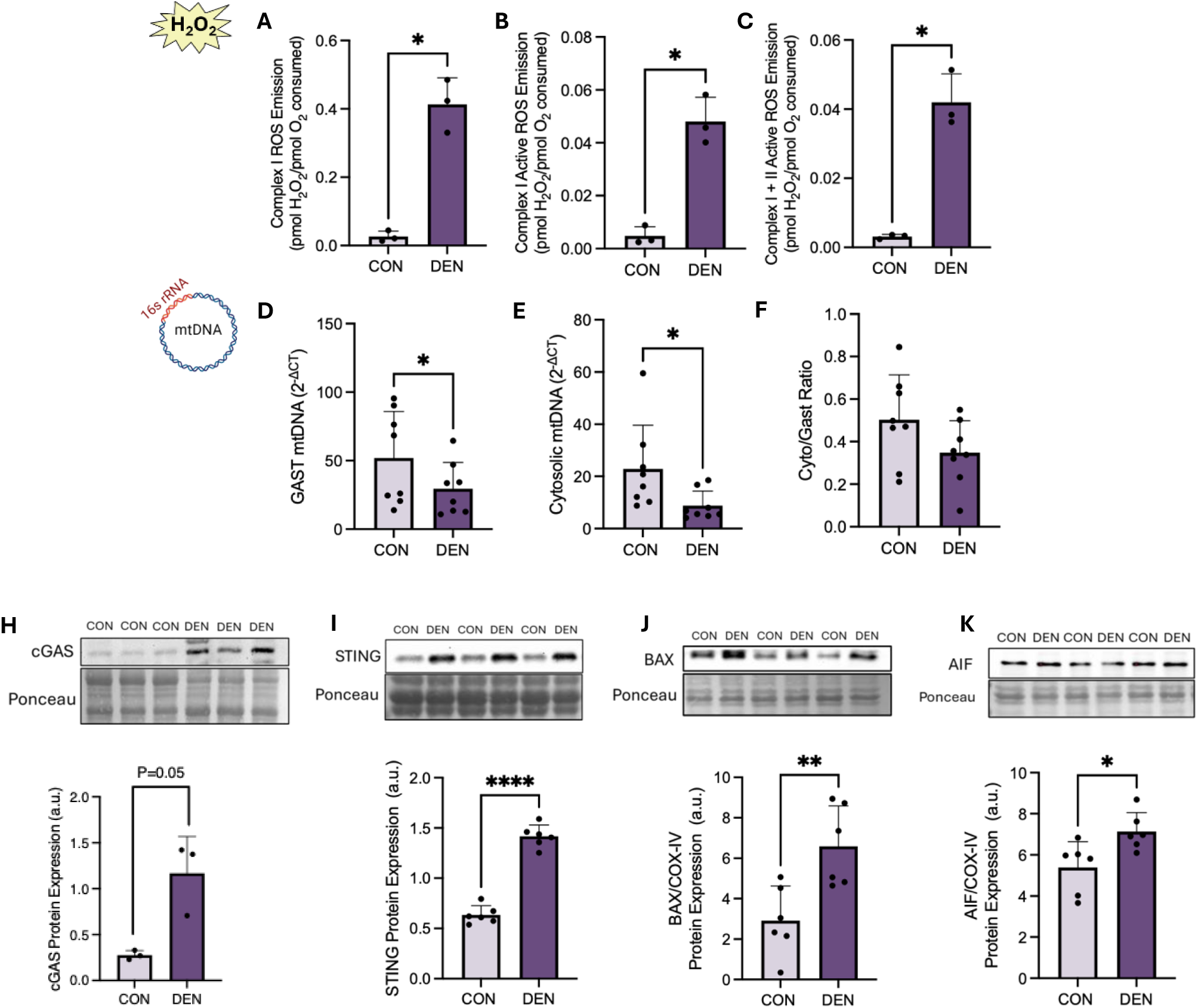
Assessment of potential DAMPs and alternative pathways following muscle denervation. ROS emission under complex I (A), complex I active (B), and complex I+II active (C) respiration states (n=3). qPCR analyses of the 16s rRNA mtDNA gene within whole muscle (D), cytosol (E), and the ratio of cytosolic/whole muscle mtDNA (F) (n=8). Representative western blots of cGAS (H, n=3), STING (I), Bax (J), and AIF (K), alongside the corresponding graphical quantifications (n=6). Data was analyzed using a paired student’s t-test. *, p<0.05; **, p<0.01; ****, p<0.0001. CON, control; DEN, denervated; Cyto, cytosolic; Gast, Gastrocnemius; All values are expressed as mean ± SD.

### Fiber-type differences of pro-inflammatory proteins in young and aged skeletal muscle

To assess the influence of skeletal muscle fiber types on the protein expression of various proteins involved in innate immune signaling, the mixed fast-twitch tibialis anterior (TA) and the slow-twitch soleus (SOL) muscles were compared in muscle from both young and aged animals. As shown in Figure 5 A, B, D-G, western blotting that the soleus muscle possessed higher levels of NLRP3, procaspase-1, STING and caspase-1 compared to the predominantly fast-twitch TA muscle, in both young and old muscle. All of these proteins were also significantly (p<0.01) elevated with age. In addition, we also observed a trending decline in COX-IV protein in aged muscle relative to young tissue, as expected.

**Figure 5:**
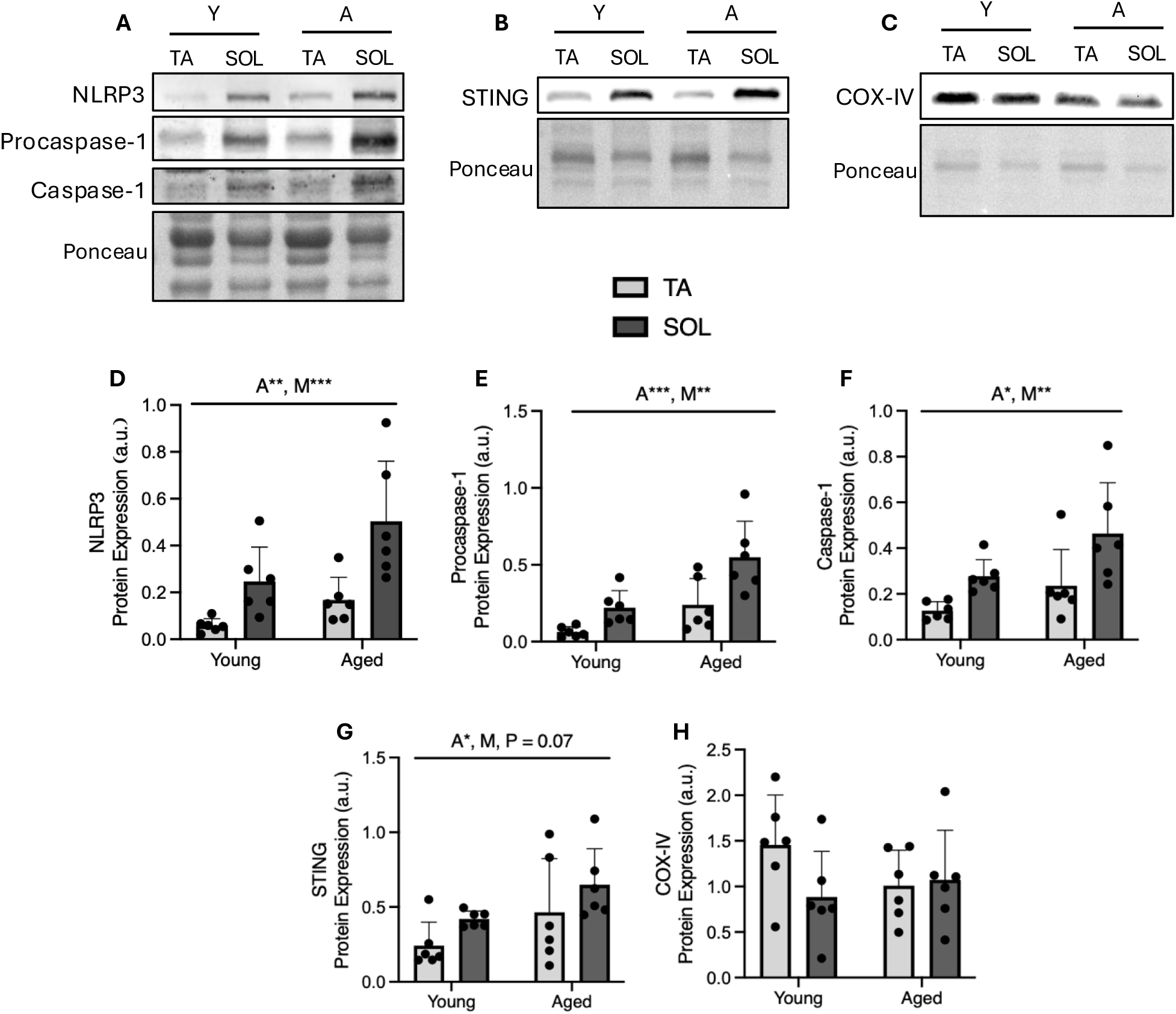
Effect of skeletal muscle fiber-type on inflammatory protein expression in young and aged skeletal muscle. Representative western blots of NLRP3 inflammasome components (A), STING protein (B), and mitochondrial content marker COX-IV (C) alongside their corresponding graphical representations (D, E, F, G, H) (n=6). To assess fiber-type differences, the tibialis anterior and soleus were used as a mixed and oxidative skeletal muscle, respectively. Data was analyzed using a two-way ANOVA. A, main effect of age; M, main effect of muscle type. *, p<0.05; **, p<0.01; ***, p<0.001. Y, young; A, aged; TA, tibialis anterior; SOL, soleus. Values are expressed as mean ± SD.

### The aging phenotype exhibits decreased muscle mass, which is recoverable with training, despite reduced running distance

After obtaining promising results demonstrating NLRP3 inflammasome activation in denervated muscle, we investigated the natural deterioration of skeletal muscle using aged mice, with or without the superimposed intervention of an endurance training protocol (Figure 6A).

**Figure 6:**
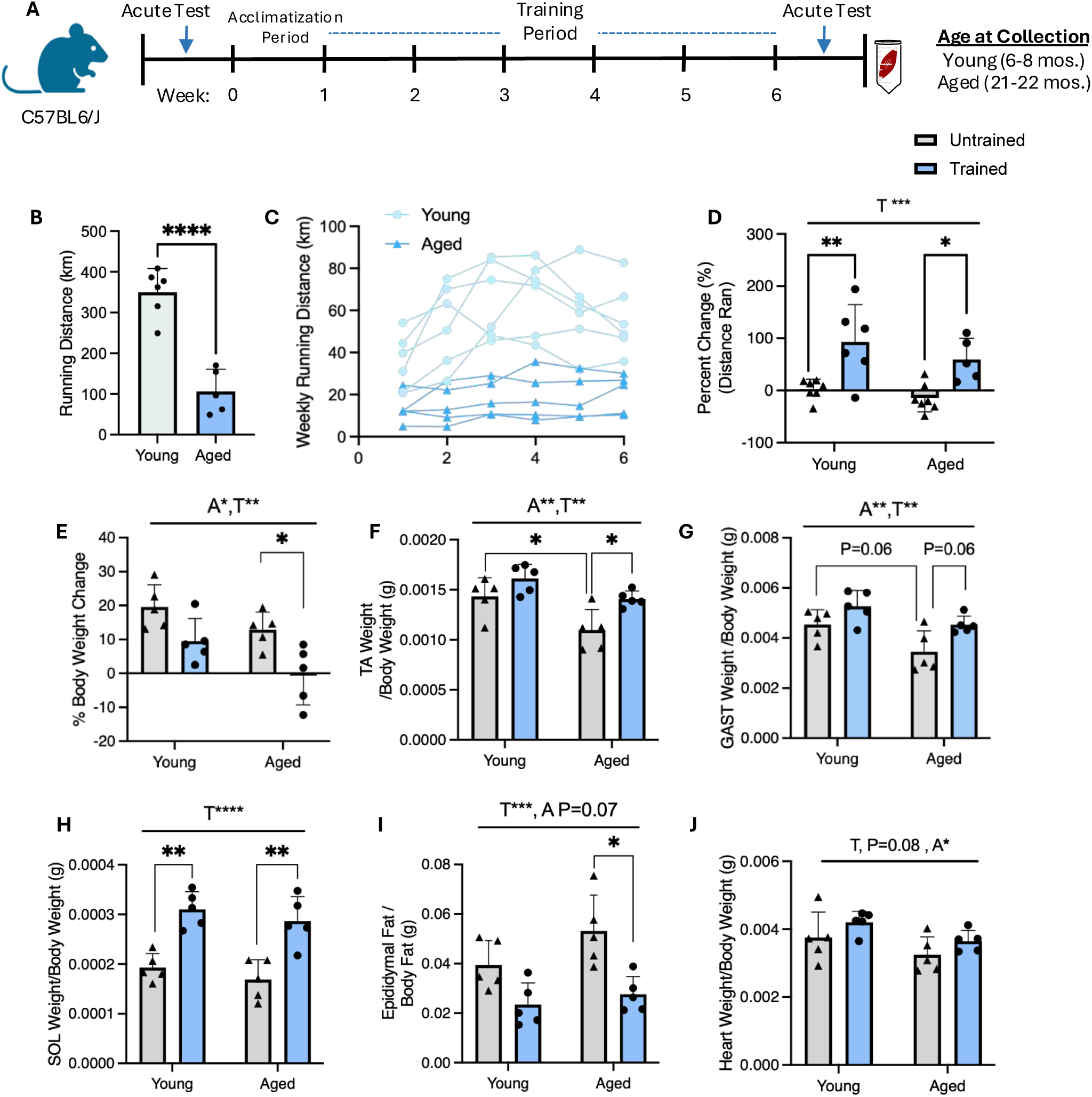
Physical characteristics in young and aged mice following 6 weeks of voluntary wheel running. Study design (A). Total distance ran over the 6-week training period for young and aged animals (km) (B) (n=5-6). Weekly running distance plotted per week (km) (C). % improvement in the distance ran during an exhaustive acute treadmill bout before and after the 6-week training protocol within young and aged animals (D) (n=5-7). % body weight change following the intervention in young and aged animals with and without training (E) (n=5). Individual muscle weights of TA (F), GAST (G) and SOL (H) corrected to body weight (n=5). Epididymal fat measurements compared across groups corrected to total body weight (I). Heart weight measurements between groups corrected to body weight (J) (n=5). Running differences were compared using an unpaired student’s t-test. Body composition was compared using a two-way ANOVA. A, main effect of age; T, main effect of training. *, p<0.05; **, p<0.01; ***, p<0.001; ****, p<0.0001. TA, tibialis anterior; SOL, soleus; GAST, gastrocnemius. Values are expressed as mean ± SD. A portion of Figure 6A was created using BioRender.com.

Throughout the training period, the aged animals displayed a reduced capacity for endurance activity as shown by a lower total running distance compared to their young counterparts (Figure 6B, 6C, p<0.0001). To ensure that both groups still demonstrated training adaptations, we assessed several physical characteristics of skeletal muscle. As shown in Figure 6E, the change in body weight taken pre- and post-intervention demonstrated a main effect of age and training, with a significant effect of training within the aged group (p<0.05). We also assessed the changes in muscle weights in the young and aged groups with training. Consistent across various mixed and highly oxidative fiber types including the tibialis anterior (Figure 6F), gastrocnemius (Figure 6G), and soleus (Figure 6H), there was a main effect of age and training, indicating reduced muscle mass basally with age, which was improved in both age groups following training. Similarly, to gain an understanding of body composition we measured epididymal fat, which demonstrated a main effect of training, especially within the trained aged mice in comparison to their untrained counterparts (Figure 6I, p<0.05). In addition, a trending main effect of training (p=0.08) and age (p<0.05) was also observed when measuring heart weight, which was used as an additional endurance training adaptation (Figure 6J). To assess a functional parameter of endurance capacity, we utilized an acute treadmill test that was performed before and after the 6-week training period. As shown in Figure 6D, the young and aged mice within the trained group showed significant improvements in their endurance capacity compared to the sedentary controls, which showed minimal adaptations.

### Aging exhibits altered mitochondrial quality, which is improved with exercise training

To assess changes in muscle mitochondria, we measured the organelle content markers COX-IV and COX-I, as subunits of complex IV in the electron transport chain. COX-IV and COX-I protein were increased with age (Figure 7B, p<0.05 and Figure 7C, p<0.01, respectively). Measures of oxygen consumption within permeabilized muscle fibers were not markedly affected by age, but we observed main effect of training in all respiration states (Figure 7D, 7E, 7F, p<0.01, p<0.05), which was more prominent in the aged animals following training (Figure 7E, p<0.05; Figure 7F, p<0.05).

**Figure 7:**
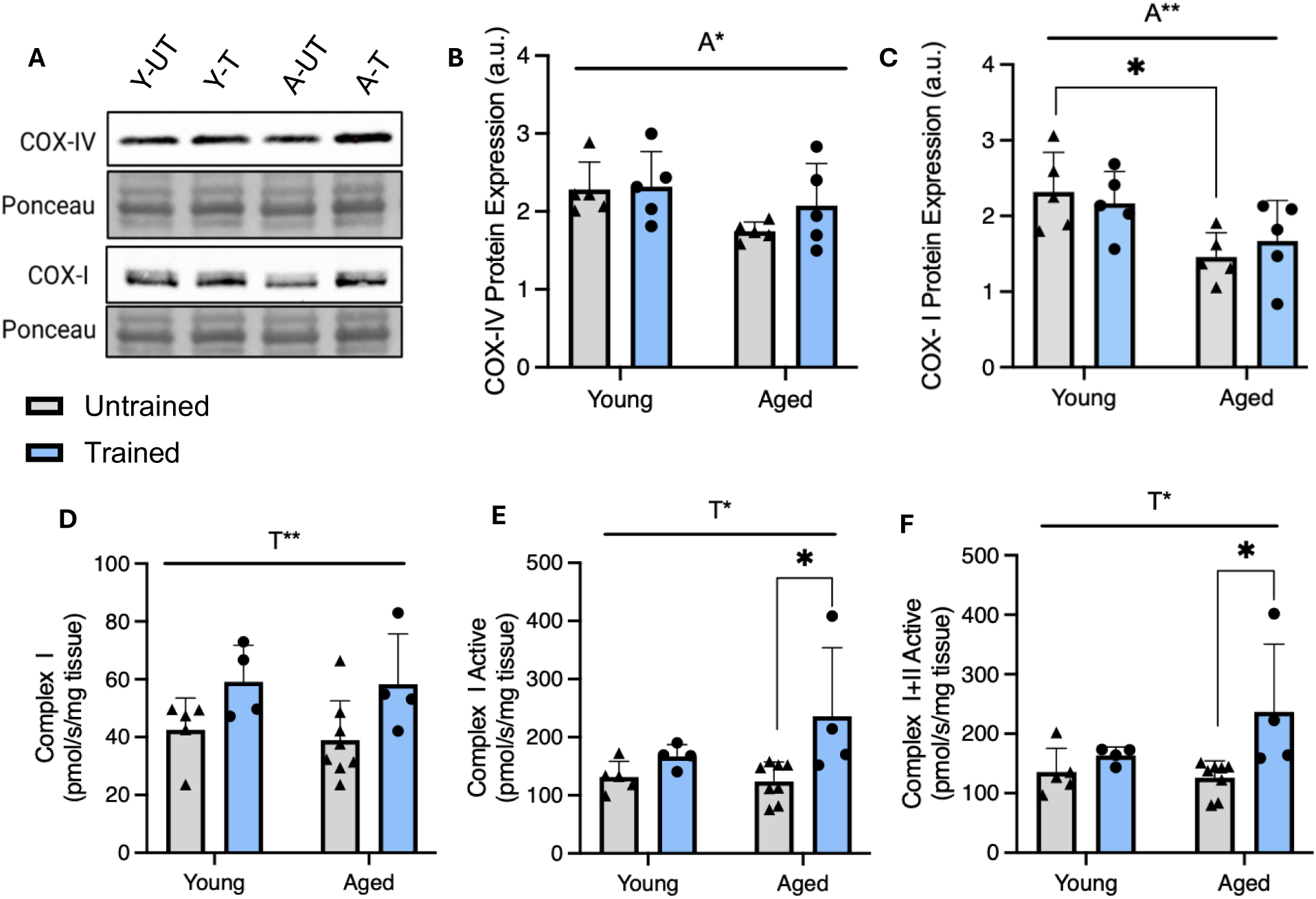
Mitochondrial adaptations in young and aged mice following the 6-week training period. Representative western blots of mitochondrial content markers COX-IV and COX-I (A), alongside the corresponding quantifications (B,C) (n=5). Oxygen consumption rates during complex I (D), complex I active (E), and complex I+II active (F) states of respiration (n=4-8). All values were analyzed using a two-way ANOVA. A, main effect of age; T, main effect of training. *, p<0.05; **, p<0.01. Values are expressed as mean ± SD.

### Aged skeletal muscle demonstrates an increased pro-inflammatory environment, which is attenuated with exercise training

After confirming that the aged animals underwent mitochondrial and skeletal muscle adaptations following exercise training, we assessed the effect of training on the innate inflammatory environment within the skeletal muscle by measuring markers of NLRP3 inflammasome activation. As an upstream measure of transcriptional activation, we observed no significant differences in the protein expression of NF-κB subunit p65 across any of the groups (Figure 8B). However, when looking at the structural components of the NLRP3 inflammasome complex, there was a main effect of age (p<0.01) and training (p<0.01) on NLRP3 protein expression, as well as a significant interaction between the two factors (p<0.01, Figure 8C). Downstream of NLRP3 inflammasome activation, we observed an increase in caspase-1 p20 protein expression with age (p<0.05; Figure 8D). Similarly, active caspase-1 protein expression exhibited a trending reduction with training following the 6-week intervention (p=0.07, Figure 8D). In addition, immature GSDMD and mature GSDMD-N showed a main effect of age (p<0.01 and p<0.05, respectively) indicating a greater accumulation of these pro-inflammatory markers (Figure 8F, 8G). However, while GSDMD protein expression decreased in aged muscle following training (p<0.05), there was no clear change in GSDMD-N (Figure 8F). When we corrected the degree of training adaptations in NLRP3 and caspase-1 to the total distance run by the young and aged animals we observed a remarkably greater attenuation of inflammatory markers compared to the young counterparts, despite the >3-fold lower exercise distance over the 6 week training period (Figure 8H, Figure 8I, p<0.05).

**Figure 8:**
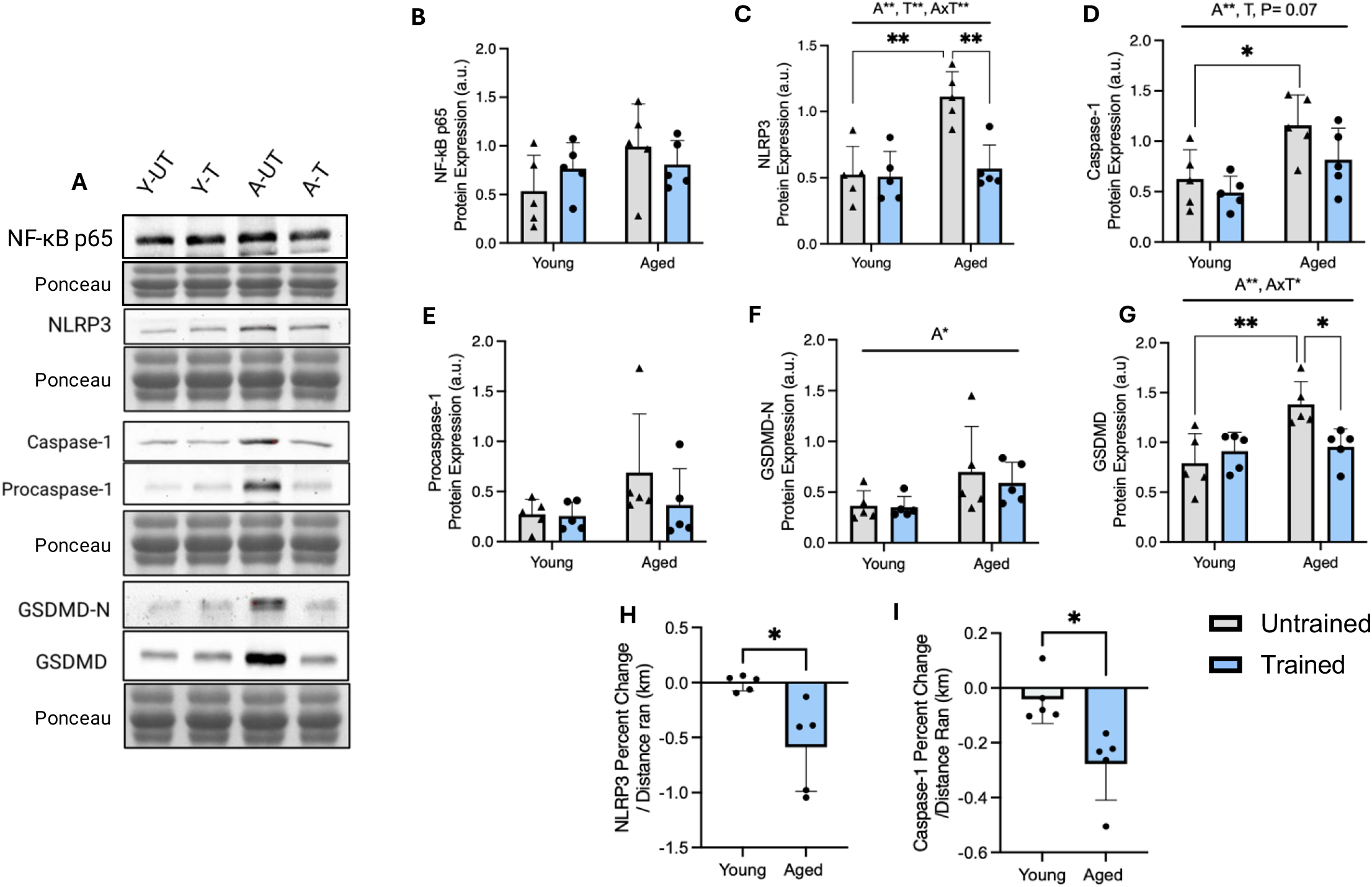
NLRP3 Inflammasome markers in young and aged animals following voluntary wheel running. Representative western blot of proteins involved in the NLRP3 Inflammasome signaling pathway (A), including NF-κB p65 (B), NLRP3 (C), Caspase-1 p20 (D), Procaspase-1 (E), GSDMD-N (F), GSDMD (G) (n=5). Fold change of NLRP3 (H) and Caspase-1 p20 (I) protein normalized to total distance ran in young and aged mice (n=5). Fold change was analyzed using an unpaired student’s t-test, while protein expression underwent a two-way ANOVA. A, main effect of age; T, main effect of training; AxT, interaction between age and training. *, p<0.05; **, p<0.01. All values are expressed as mean ± SD.

### Age and training demonstrated no effect on ROS and mtDNA release but showed increased expression of innate immune markers and mPTP components

After observing the apparent effects of both age and training on NLRP3 inflammasome activation, we wanted to further characterize the presence of DAMPS within whole muscle and cytosolic fractions. With both age and training, there were no apparent changes in H_2_O_2_ production following complex I inactive (Figure 9A), complex I active (Figure 9B), and complex I+II active respiration (Figure 9C). When investigating the presence of mtDNA as a mtDAMP, there were no significant changes in whole muscle mtDNA and cytosolic mtDNA following the intervention (Figure 9D, 9E). However, the amount of cytosolic mtDNA relative to whole muscle mtDNA was increased with age and attenuated by training (Figure 9F, p<0.05), indicating a differential response to endurance training in aged and young animals. This related well to changes in the upstream release protein Bax, which was upregulated basally within aging muscle (Figure 9I, p<0.01) and attenuated with training (p<0.05). In addition, it matched closely with the changes in downstream STING expression, which exhibited a trending reduction with training, an effect that was most apparent within the aged muscle (Figure 9G, p=0.05). In addition, aged muscle demonstrated a larger capacity to attenuate STING protein expression with reduced running distance, similar to NLRP3 and caspase-1 (Figure 9H, p<0.05).

**Figure 9:**
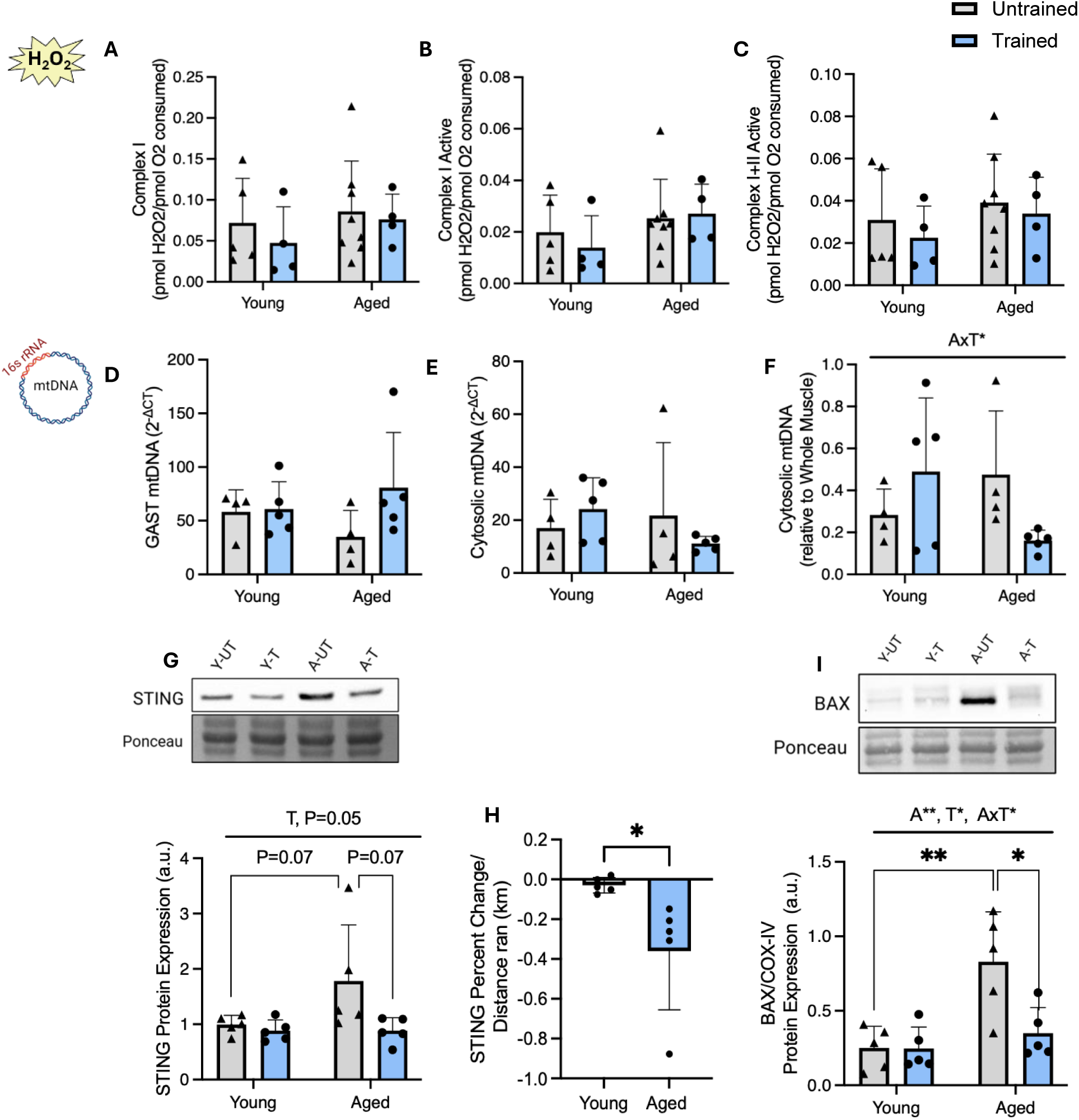
Analysis of potential DAMPS and associated pathways in young and aged animals following voluntary wheel running. ROS emission under complex I (A), complex I active (B), and complex I+II active (C) respiration states (n=4-8). qPCR analyses of the 16s rRNA mtDNA fragment within whole muscle (D), cytosol (E), and the ratio of cytosolic/whole muscle mtDNA (F) (n=4-5). Representative western blots of STING (G) and BAX (I) protein alongside the corresponding graphical quantifications (n=5). Fold change of STING protein (H) corrected to total distance ran in young and aged mice (n=5). Data was analyzed using a two-way ANOVA. A, main effect of age; T, main effect of training; AxT, interaction between age and training; *, p<0.05; **, p<0.01; Cyto, cytosolic; Gast, Gastrocnemius; All values are expressed as mean ± SD.

**Figure 10:**
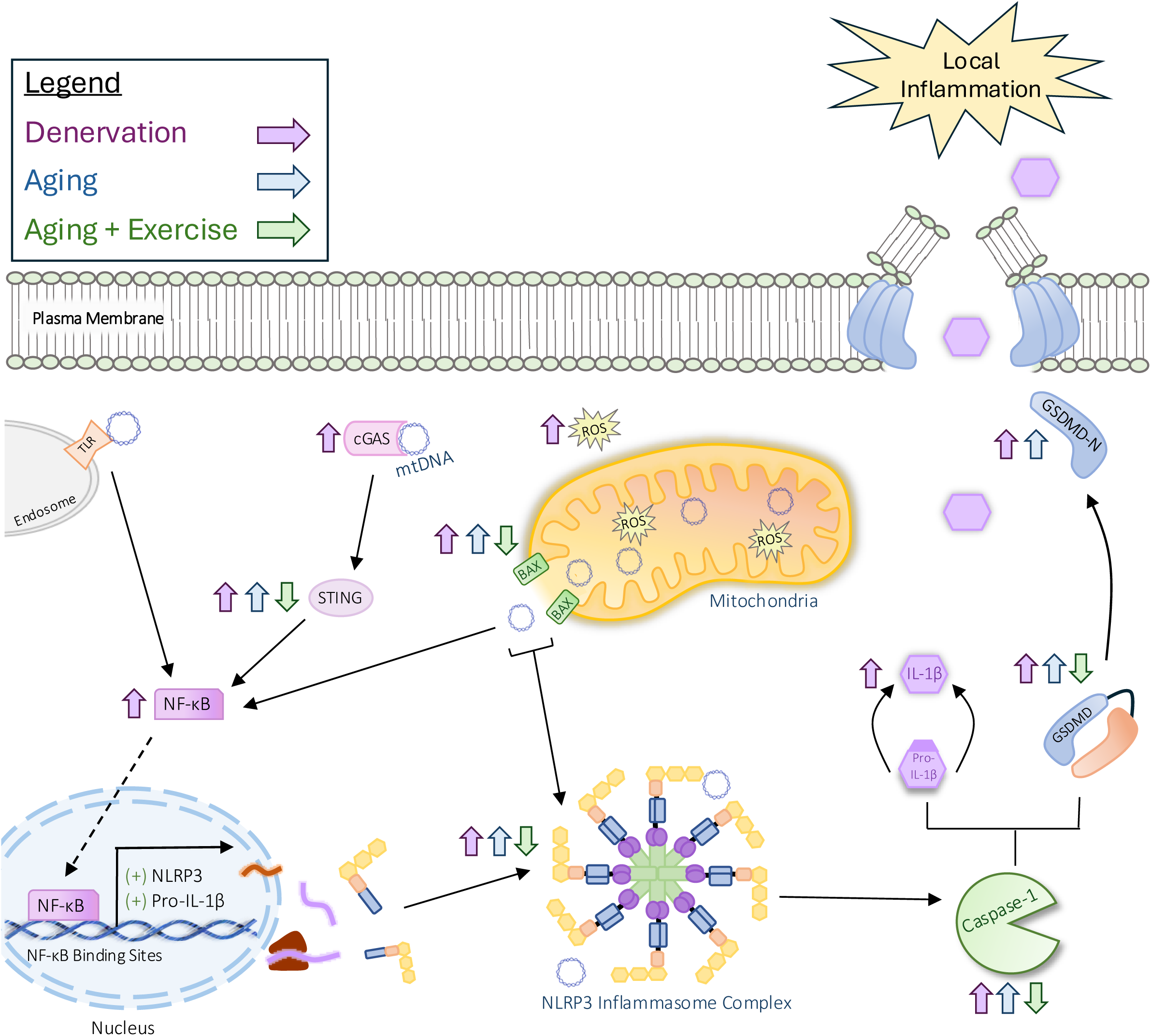
Summary of the effects of muscle denervation, age, and endurance training on the expression on innate immune signaling proteins. Various mitochondrial DAMPs can augment the activation of the NLRP3 inflammasome complex to promote its assembly and the subsequent maturation of caspase-1, GSDMD-N and IL-1β. Following 7 days of muscle denervation and with age, there were observable increases in the protein expression of cGAS-STING and NF-κB, and BAX which partially mediates the mitochondrial permeability transition pore. Furthermore, when assessing downstream components of the NLRP3 inflammasome, several proteins such as NLRP3, caspase-1, GSDMD, IL-1β, and GSDMD-N were increased, indicating a pro-inflammatory environment. When investigating the combined effect of aging and chronic endurance training, several proteins that have been implicated in innate immune signaling including BAX, STING, NLRP3, caspase-1 and GSDMD demonstrated an attenuation of the basal increase that was observed.

## Discussion

The purpose of this study was to investigate *in vivo* innate immune signaling, specifically focused on the NLRP3 inflammasome complex in response to skeletal muscle use, disuse, and aging. Mitochondria are now being explored as critical signaling hubs that can mediate inflammatory processes, however, the relationship between mitochondrial health, skeletal muscle activity, and innate immune activation requires additional work. As a result, we utilized a potent model of muscle disuse through a sciatic nerve transection to assess the response of the NLRP3 inflammasome complex. Furthermore, aging has been established as a complex condition associated with skeletal muscle deterioration, inflammation, and functional detriments which prompted us to investigate the downstream consequences and potential benefits of endurance training. By assessing mitochondrial parameters and NLRP3 inflammasome activation, we aim to understand the inflammatory environment in relation to skeletal muscle mitochondrial adaptations.

Consistent with other findings, 7 days of muscle denervation upregulated catabolic signaling pathways as shown through an increased expression of the E3 ubiquitin ligase MURF1 and muscle atrophy marker GADD45-α, leading to a notable loss of muscle mass (18). Furthermore, we observed consistent decreases in mitochondrial content and function, which has also been shown in several other studies utilizing a similar model (10, 19). This suggests that 7 days of muscle denervation is sufficient to induce maladaptive changes within the mitochondrial pool, which encouraged us to investigate changes in NLRP3 inflammasome signaling. Previous work by You and colleagues demonstrated a relationship between denervation-induced atrophy and upregulation of proteins involved in the NLRP3 signaling cascade including NLRP3, caspase-1, GSDMD-N, and IL-1β following 14-28 days of denervation, which was attenuated by NLRP3 KO (11). This relationship was further demonstrated as LPS-induced NLRP3 inflammasome activation led to decreases in myotube diameter that was rescued upon treatment with the NLRP3 inhibitor, MCC950 (12). In our study, we demonstrated an early upregulation in the protein levels of NF-κB p65, NLRP3, procaspase-1, caspase-1 p20, GSDMD, GSDMD-N, and IL-1β as soon as 7 days following the sciatic nerve transection, suggesting that NLRP3 inflammasome activation occurs at earlier time points of denervation-induced muscle disuse than previously described.

Since it is known that mtDAMPS can act as potent activators of the NLRP3 inflammasome (20, 21), we measured the production of ROS and mtDNA release to understand potential upstream activators. Consistent with other findings, 7 days of denervation induced dramatic increases in the production of ROS which could be contributing to the observed inflammasome activation (10, 20, 22). In addition, the release of mtDNA into the cytosol has been widely recognized as a pro-inflammatory molecule, which encouraged us to use a mtDNA-specific 16s rRNA primer to assess its localization. Previous research has demonstrated that maladaptive changes in mitochondria such as impairments in autophagy (23) or mitochondrial dynamics (24) have been associated with the release of mtDNA and subsequent upregulation of NLRP3 inflammasome signaling. In contrast to our original hypothesis, the amount of mtDNA within the cytosolic relative to total mtDNA was not altered with 7 days of denervation. However, marked elevations in the expression of cGAS and STING protein, which are involved in the recognition of double-stranded DNA within the cytosol were observed, indicating that an increased sensitivity to mtDNA toward inflammasome activation likely exists with denervation. Furthermore, several studies have specifically implicated oxidized mtDNA in its ability to bind to and stimulate NLRP3 inflammasome assembly (21, 25). While we observed no changes in the quantity of cytosolic mtDNA relative to whole muscle, the elevated oxidative stress could contribute to a higher proportion of oxidized mtDNA within denervated muscle, which has yet to be explored. In addition, others have demonstrated that cGAS-STING pathway activation can promote the nuclear translocation of NF-κB, and converge on NLRP3 inflammasome activation in a STING-dependent manner (6). Several groups have also implicated the formation of the mitochondrial permeability transition pore in augmenting the release of mtDAMPS and activation of innate immune responses, which is supported by the increased BAX and AIF protein expression that we observed following muscle denervation (25, 26). Finally, as we noted increases in NLRP3 inflammasome activation as early as seven days post-denervation, this could suggest a need to assess earlier time points than 7 days to identify the upstream signals involved.

Similarly, the aging phenotype is associated with chronic low-grade inflammation and deteriorations in mitochondrial function, muscle mass, and functional performance, prompting us to investigate NLRP3 inflammasome expression and mitochondrial parameters (27). Potential interventions such as exercise are in high demand to address the skeletal muscle deterioration associated with aging. While the aged animals displayed a reduced willingness to train compared to their young counterparts, they still exhibited a variety of training-induced adaptations including increases in heart mass and hindlimb muscle mass, consistent with other studies employing the same model (28, 29). The trained young and aged mice also demonstrated improved body composition evident through a reduction of fat mass and reduced weight gain in comparison to the untrained controls. Aged animals displayed a reduced mitochondrial content, which is also considered a consistent characteristic of aging skeletal muscle and could contribute to their reduced running capacity (30). However, the training period was still able to induce an increase oxygen consumption, particularly within the aged animals, demonstrating that the intervention was sufficient to enhance mitochondrial function, despite different running capacities.

As with the denervation study, we investigated the innate inflammatory profile within skeletal muscle to investigate the relationship between mitochondrial function and NLRP3 inflammasome activation. We observed a consistent elevation of the inflammatory markers STING, NLRP3, procaspase-1, caspase-1, GSDMD, and GSDMD-N within aged skeletal muscle. Others have shown comparable findings associated with increased muscle atrophy and caspase-1 activity in 24-month old mice (14). Our results illustrate a novel beneficial role of endurance training in attenuating the age-related inflammatory profile within aged skeletal muscle, as the training intervention was able to reduce the protein expression of NLRP3, caspase-1 and GSDMD, as well as attenuate the levels of cytosolic mtDNA in aged animals. As NLRP3 inflammasome activity has been linked to muscle atrophy within muscle, this anti-inflammatory effect of exercise could be contributing to the restoration of muscle mass observed in the trained aged animals. This aligns with other studies that have demonstrated reduced TLR/NF-κB signaling and NLRP3 inflammasome activation in hippocampal cells and adipose tissue following long-term treadmill training in conditions such as Parkinson’s disease and diet-induced obesity (16, 31). Furthermore, aging was also associated with an increased expression of STING and BAX, similarly to denervation, which could further augment the inflammatory profile within muscle via the release of mtDAMPS. Most importantly, following the same trend that was observed with muscle mass and mitochondrial function adaptations, aged skeletal muscle displayed a larger attenuation of inflammatory proteins, including STING and BAX, despite a lower exercise capacity, suggesting that innate immune signaling in aged muscle is increasingly responsive to endurance exercise compared to young, healthy controls. Thus, while the absolute quantity of exercise performed by the aged animals was reduced compared to the young mice, the relative workload was still sufficient to induce beneficial adaptations in mitochondrial function and innate immune protein expression. Within the literature, there is still conflicting evidence surrounding the adaptive potential of aging skeletal muscle. While some groups have demonstrated that aging muscle possesses the same capacity for improvement as young muscle (32), other work has shown an attenuated reduced adaptive response specifically targeted to mitochondrial changes with age (33). However, consistent with our results, exercise has been shown to induce greater reductions in pro-apoptotic signaling in aged muscle (33).

In addition to the basal increase in pro-inflammatory proteins with age, our data demonstrates the novel finding of fiber-type differences within skeletal muscle. When compared to the fast-twitch mixed TA, our results show higher basal levels of the pro-inflammatory proteins NLRP3, procaspase-1, caspase-1 and STING protein expression in the slow-twitch soleus muscle, which persisted even in the aged cohort. Supporting this, Zimowska et al. showed that following acute muscle injury, the inflammatory response of the soleus muscle was more potent and prolonged compared to the fast-twitch EDL when assessing immune cell infiltration and cytokine expression (34). Further work on the trained immune responses within different muscle fiber types is warranted.

In summary, our study demonstrates that skeletal muscle disuse promotes NLRP3 inflammasome activation and mitochondrial deterioration, contributing to a pro-inflammatory environment within skeletal muscle. This work emphasizes the anti-inflammatory properties of chronic endurance training in attenuating the age-related increases in NLRP3 inflammasome activation and improving mitochondrial function, and more work is required 1) to identify the regulatory pathway that leads to the exercise-induced down-regulation of pro-inflammatory genes in aged muscle, and 2) to investigate the presence of sex differences in innate immune adaptations to exercise as there is sufficient evidence to suggest sexual dimorphisms in mitochondrial and innate immune adaptations (35, 36). Surprisingly, these results also suggest an increased adaptive response of NLRP3 inflammasome activation to endurance training within aged skeletal muscle, even in the presence of a lower exercise quantity compared to the young mice. This is consistent with the general, emerging idea that even smaller doses of exercise can be beneficial (37), especially in the aging population. As overstimulated NLRP3 inflammasome activation is associated with a variety of disease states, this work helps establish exercise training as a promising intervention for combatting inflammation within skeletal muscle. Furthermore, this also emphasizes that an exercise intolerant aged population can still reap the anti-inflammatory benefits of endurance training at lower intensities, making this a more feasible intervention.

## Manuscript Author Contributions

**Priyanka Khemraj:** Performed denervation surgeries and tissue collection for all animals; collected data for chronic and acute training protocols; conducted experimental testing and data analysis for DNA extractions, subcellular isolations, qPCR, ELISA, respiration and ROS analysis, and western blotting; wrote the manuscript.

**Anastasiya Kuznyetsova:** Aided in performing mitochondrial fractionations and western blotting.

**Dr. David A. Hood:** Supervisor of the project and is the principal investigator.

## Acknowledgments

This work was supported by funding from the Natural Sciences and Engineering Research Council of Canada (NSERC) to D.A. Hood. P. Khemraj is a recipient of the Ontario Graduate Scholarship.

## References

1. Brunelli S, Roverequerini P. The immune system and the repair of skeletal muscle. Pharmacological Research 58: 117–121, 2008. doi: 10.1016/j.phrs.2008.06.008.

2. Chen Y-F, Lee C-W, Wu H-H, Lin W-T, Lee OK. Immunometabolism of macrophages regulates skeletal muscle regeneration. Front Cell Dev Biol 10: 948819, 2022. doi: 10.3389/fcell.2022.948819.

3. De Paepe B, De Bleecker JL. Cytokines and Chemokines as Regulators of Skeletal Muscle Inflammation: Presenting the Case of Duchenne Muscular Dystrophy. Mediators of Inflammation 2013: 1–10, 2013. doi: 10.1155/2013/540370.

4. Loell I, Lundberg IE. Can muscle regeneration fail in chronic inflammation: a weakness in inflammatory myopathies?: Review: Muscle weakness in myositis. Journal of Internal Medicine 269: 243–257, 2011. doi: 10.1111/j.1365-2796.2010.02334.x.

5. Kelley N, Jeltema D, Duan Y, He Y. The NLRP3 Inflammasome: An Overview of Mechanisms of Activation and Regulation. IJMS 20: 3328, 2019. doi: 10.3390/ijms20133328.

6. Zhang W, Li G, Luo R, Lei J, Song Y, Wang B, Ma L, Liao Z, Ke W, Liu H, Hua W, Zhao K, Feng X, Wu X, Zhang Y, Wang K, Yang C. Cytosolic escape of mitochondrial DNA triggers cGAS-STING-NLRP3 axis-dependent nucleus pulposus cell pyroptosis. Exp Mol Med 54: 129–142, 2022. doi: 10.1038/s12276-022-00729-9.

7. Li D, Wu M. Pattern recognition receptors in health and diseases. Sig Transduct Target Ther 6: 291, 2021. doi: 10.1038/s41392-021-00687-0.

8. He W, Wan H, Hu L, Chen P, Wang X, Huang Z, Yang Z-H, Zhong C-Q, Han J. Gasdermin D is an executor of pyroptosis and required for interleukin-1β secretion. Cell Res 25: 1285–1298, 2015. doi: 10.1038/cr.2015.139.

9. Gurung P, Lukens JR, Kanneganti T-D. Mitochondria: diversity in the regulation of the NLRP3 inflammasome. Trends in Molecular Medicine 21: 193–201, 2015. doi: 10.1016/j.molmed.2014.11.008.

10. Triolo M, Slavin M, Moradi N, Hood DA. Time□dependent changes in autophagy, mitophagy and lysosomes in skeletal muscle during denervation□induced disuse. The Journal of Physiology 600: 1683– 1701, 2022. doi: 10.1113/JP282173.

11. You Z, Huang X, Xiang Y, Dai J, Xu L, Jiang J, Xu J. Ablation of NLRP3 inflammasome attenuates muscle atrophy via inhibiting pyroptosis, proteolysis and apoptosis following denervation. Theranostics 13: 374–390, 2023. doi: 10.7150/thno.74831.

12. Eggelbusch M, Shi A, Broeksma BC, Vázquez-Cruz M, Soares MN, De Wit GMJ, Everts B, Jaspers RT, Wüst RCI. The NLRP3 inflammasome contributes to inflammation-induced morphological and metabolic alterations in skeletal muscle. J cachexia sarcopenia muscle 13: 3048–3061, 2022. doi: 10.1002/jcsm.13062.

13. Franceschi C, Bonafè M, Valensin S, Olivieri F, De Luca M, Ottaviani E, De Benedictis G. Inflamm-aging: An Evolutionary Perspective on Immunosenescence. Annals of the New York Academy of Sciences 908: 244–254, 2000. doi: 10.1111/j.1749-6632.2000.tb06651.x.

14. McBride MJ, Foley KP, D’Souza DM, Li YE, Lau TC, Hawke TJ, Schertzer JD. The NLRP3 inflammasome contributes to sarcopenia and lower muscle glycolytic potential in old mice. American Journal of Physiology-Endocrinology and Metabolism 313: E222–E232, 2017. doi: 10.1152/ajpendo.00060.2017.

15. Chen CCW, Erlich AT, Hood DA. Role of Parkin and endurance training on mitochondrial turnover in skeletal muscle. Skeletal Muscle 8: 10, 2018. doi: 10.1186/s13395-018-0157-y.

16. Wang W, Lv Z, Gao J, Liu M, Wang Y, Tang C, Xiang J. Treadmill exercise alleviates neuronal damage by suppressing NLRP3 inflammasome and microglial activation in the MPTP mouse model of Parkinson’s disease. Brain Res Bull 174: 349–358, 2021. doi: 10.1016/j.brainresbull.2021.06.024.

17. Lee J, Lee Y, LaVoy EC, Umetani M, Hong J, Park Y. Physical activity protects NLRP3 inflammasome-associated coronary vascular dysfunction in obese mice. Physiol Rep 6: e13738, 2018. doi: 10.14814/phy2.13738.

18. Bongers KS, Fox DK, Ebert SM, Kunkel SD, Dyle MC, Bullard SA, Dierdorff JM, Adams CM. Skeletal muscle denervation causes skeletal muscle atrophy through a pathway that involves both Gadd45a and HDAC4. American Journal of Physiology-Endocrinology and Metabolism 305: E907–E915, 2013. doi: 10.1152/ajpendo.00380.2013.

19. Adhihetty PJ, O’Leary MFN, Chabi B, Wicks KL, Hood DA. Effect of denervation on mitochondrially mediated apoptosis in skeletal muscle. Journal of Applied Physiology 102: 1143–1151, 2007. doi: 10.1152/japplphysiol.00768.2006.

20. Zhou R, Yazdi AS, Menu P, Tschopp J. A role for mitochondria in NLRP3 inflammasome activation. Nature 469: 221–225, 2011. doi: 10.1038/nature09663.

21. Shimada K, Crother TR, Karlin J, Dagvadorj J, Chiba N, Chen S, Ramanujan VK, Wolf AJ, Vergnes L, Ojcius DM, Rentsendorj A, Vargas M, Guerrero C, Wang Y, Fitzgerald KA, Underhill DM, Town T, Arditi M. Oxidized mitochondrial DNA activates the NLRP3 inflammasome during apoptosis. Immunity 36: 401–414, 2012. doi: 10.1016/j.immuni.2012.01.009.

22. Russell-Guzmán J, Américo-Da Silva L, Cadagan C, Maturana M, Palomero J, Estrada M, Barrientos G, Buvinic S, Hidalgo C, Llanos P. Activation of the ROS/TXNIP/NLRP3 pathway disrupts insulin-dependent glucose uptake in skeletal muscle of insulin-resistant obese mice. Free Radic Biol Med 222: 187– 198, 2024. doi: 10.1016/j.freeradbiomed.2024.06.011.

23. Nakahira K, Haspel JA, Rathinam VAK, Lee S-J, Dolinay T, Lam HC, Englert JA, Rabinovitch M, Cernadas M, Kim HP, Fitzgerald KA, Ryter SW, Choi AMK. Autophagy proteins regulate innate immune responses by inhibiting the release of mitochondrial DNA mediated by the NALP3 inflammasome. Nat Immunol 12: 222–230, 2011. doi: 10.1038/ni.1980.

24. Irazoki A, Gordaliza-Alaguero I, Frank E, Giakoumakis NN, Seco J, Palacín M, Gumà A, Sylow L, Sebastián D, Zorzano A. Disruption of mitochondrial dynamics triggers muscle inflammation through interorganellar contacts and mitochondrial DNA mislocation. Nat Commun 14: 108, 2023. doi: 10.1038/s41467-022-35732-1.

25. Xian H, Watari K, Sanchez-Lopez E, Offenberger J, Onyuru J, Sampath H, Ying W, Hoffman HM, Shadel GS, Karin M. Oxidized DNA fragments exit mitochondria via mPTP- and VDAC-dependent channels to activate NLRP3 inflammasome and interferon signaling. Immunity 55: 1370–1385.e8, 2022. doi: 10.1016/j.immuni.2022.06.007.

26. Yamazaki T, Galluzzi L. BAX and BAK dynamics control mitochondrial DNA release during apoptosis. Cell Death Differ 29: 1296–1298, 2022. doi: 10.1038/s41418-022-00985-2.

27. López-Otín C, Blasco MA, Partridge L, Serrano M, Kroemer G. The Hallmarks of Aging. Cell 153: 1194–1217, 2013. doi: 10.1016/j.cell.2013.05.039.

28. Wong JC, Oliveira AN, Khemraj P, Hood DA. The role of TFE3 in mediating skeletal muscle mitochondrial adaptations to exercise training. Journal of Applied Physiology 136: 262–273, 2024. doi: 10.1152/japplphysiol.00484.2023.

29. Crilly MJ, Tryon LD, Erlich AT, Hood DA. The role of Nrf2 in skeletal muscle contractile and mitochondrial function. J Appl Physiol (1985) 121: 730–740, 2016. doi: 10.1152/japplphysiol.00042.2016.

30. Chabi B, Ljubicic V, Menzies KJ, Huang JH, Saleem A, Hood DA. Mitochondrial function and apoptotic susceptibility in aging skeletal muscle. Aging Cell 7: 2–12, 2008. doi: 10.1111/j.1474-9726.2007.00347.x.

31. Mardare C, Krüger K, Liebisch G, Seimetz M, Couturier A, Ringseis R, Wilhelm J, Weissmann N, Eder K, Mooren F-C. Endurance and Resistance Training Affect High Fat Diet-Induced Increase of Ceramides, Inflammasome Expression, and Systemic Inflammation in Mice. J Diabetes Res 2016: 4536470, 2016. doi: 10.1155/2016/4536470.

32. Škorjanc D, Traub I, Pette D. Identical responses of fast muscle to sustained activity by low-frequency stimulation in young and aging rats. Journal of Applied Physiology 85: 437–441, 1998. doi: 10.1152/jappl.1998.85.2.437.

33. Ljubicic V, Joseph A-M, Adhihetty PJ, Huang JH, Saleem A, Uguccioni G, Hood DA. Molecular basis for an attenuated mitochondrial adaptive plasticity in aged skeletal muscle. Aging 1: 818–830, 2009. doi: 10.18632/aging.100083.

34. Zimowska M, Kasprzycka P, Bocian K, Delaney K, Jung P, Kuchcinska K, Kaczmarska K, Gladysz D, Streminska W, Ciemerych MA. Inflammatory response during slow- and fast-twitch muscle regeneration: Inflammation and Muscle Repair. Muscle Nerve 55: 400–409, 2017. doi: 10.1002/mus.25246.

35. Oliveira AN, Memme JM, Wong J, Hood DA. Dimorphic effect of TFE3 in determining mitochondrial and lysosomal content in muscle following denervation. Skeletal Muscle 14: 7, 2024. doi: 10.1186/s13395-024-00339-1.

36. Jaillon S, Berthenet K, Garlanda C. Sexual Dimorphism in Innate Immunity. Clinic Rev Allerg Immunol 56: 308–321, 2019. doi: 10.1007/s12016-017-8648-x.

37. 37. Lee I-M, Powell KE, Sarmiento OL, Hallal PC. Even a small dose of physical activity can be good medicine.

